# FragLlama: Next-fragment prediction for molecular design

**DOI:** 10.1101/2024.09.28.615626

**Authors:** Jian Shen, Shengmin Zhou, Xing Che

## Abstract

The emergence of ChatGPT has drawn significant attention to Large Language Models (LLMs) due to their impressive performance. While LLMs primarily focus on next token/word prediction, we apply this principle to molecular design by reframing the task as predicting the next token/fragment. We present FragLlama, a large language model trained for molecular design, featuring custom tokens that represent molecular fragments and functional groups. The model is for generating molecules given one or two fragments, for application scenarios like general hit-to-lead and lead optimization stage drug design, PROTAC linker design; mapping to commonly used drug design strategies like fragment growing and scaffold hopping. In the pre-training stage, we adapted the Llama 3 architecture to create FragLlama, training it to learn conditional probabilities of these fragment-level tokens. The subsequent alignment stage employed fine-tuning to guide the model towards generating molecules with desired properties. The effectiveness of FragLlama is demonstrated through its applications in designing molecular glue libraries, PROTAC linkers and EGFR binders. FragLlama demonstrates proficiency in reproducing expert-level designs while also exploring novel and promising chemical spaces, highlighting its potential to augment the capabilities of medicinal chemists in drug design.

## Introduction

### What is LLM and why it’s so successful

Large Language Models (LLMs) represent a major breakthrough in language modeling, building upon decades of advancements since Shannon’s application of information theory in the 1950s.^1^ Over time, various model architectures such as Linear Models ^2^ and Convolutional Neural Networks (CNNs)^3^ emerged and dominated the field, only to reach their performance limits, necessitating new approaches. These innovations were often propelled by advances in hardware, which enabled machine learning models to leverage more complex statistical methods and larger datasets.

The success of LLMs, particularly those based on deep neural networks, can be attributed to two key principles. First, the concept of Universal Function Approximation^4,5^ suggests that large neural networks, given enough neurons and depth, can theoretically approximate any continuous function. This capability allows LLMs to model highly complex relationships in data, making them powerful tools for understanding and generating language. As deep learning progressed, this principle became the foundation for building increasingly large and sophisticated models that could capture nuanced patterns in high-dimensional data.^6^

Second, the Turing completeness of Transformer-based architectures further enhances the power of LLMs.^7,8^ While earlier models like Recurrent Neural Networks (RNNs) and Long Short-Term Memory (LSTM) networks are also Turing complete, the Transformer architecture introduces significant advantages. Transformers,^9^ through their self-attention mechanisms and parallelizable feed-forward networks, can process and comprehend vast amounts of contextual information more efficiently than their predecessors. This architectural innovation enables LLMs to perform a wide range of tasks with remarkable accuracy and flexibility.

LLMs, such as those based on the Generative Pre-trained Transformer (GPT),^10,11^ leverage these principles—Universal Function Approximation and Turing completeness— to excel in natural language processing tasks. Pre-training on massive datasets, followed by fine-tuning for specific applications, enables these models to autonomously discover and extract intricate patterns in language, resulting in performance and capabilities that far surpass those of smaller or earlier models.

### Why LLM for molecular design and the challenges

The success of LLMs in natural language processing has sparked interest in their application to other domains, including molecular design. LLMs possess two key strengths that make them well-suited for this task: representation power and generative capabilities. Their ability to capture complex patterns and relationships is critical for understanding molecular structures and properties. Additionally, LLMs’ proficiency in generating novel content aligns with the need to design new molecules. Several chemistry-specific language models have already been developed, including but not limited to ChemSpaceAL,^12^ cMolGPT,^13^ Regression Transformer,^14^ MolGPT,^15^ SAFE-GPT,^16^ and Darwin.^17^ For a more comprehensive review, please refer to the paper A Review of Large Language Models and Autonomous Agents in Chemistry.^18^

Compared to molecular representation using graph-based methods, the utilization of Transformer architectures offers superior scalability and the potential to uncover latent features that may elude human perception. Like Ilya Sutskever ever commented, ”Predicting the next token well means that you understand the underlying reality that led to the creation of that token. It’s not just statistics.”^19^ This principle is directly applicable to molecular design, where LLMs can be used to infer and predict molecular properties in a manner akin to how they process natural language.

While the application of Language Models in Chemistry presents promising avenues for research, it is imperative to acknowledge several significant challenges. This study highlights a few key issues:

1. Sequentialization of molecular information: Human languages are inherently sequential, and observations from GPT-like Large Language Models (LLMs) indicate that the token count for expressing identical information varies considerably across languages, as does the model’s logical reasoning capability. In the field of chemistry, however, there is no consensus on the optimal ”molecular language” for the sequential representation of molecules.
2. Diversity and complexity of chemical systems: The vast heterogeneity among chemical systems presents a major obstacle to developing universally applicable models for designing effective drug molecules. Additionally, the limited availability of high-quality, labeled molecular data hinders the alignment stage training, reducing the model’s ability to design molecules with specific desired properties. In general, the lack of quality data impedes the development of models at a sufficient scale to unlock emergent capabilities.
3. Evaluation challenges: Evaluating molecule language models is more complex than traditional LLMs. Validating the qualities of designed molecules often requires lengthy experimental procedures, resulting in extended feedback times.

These challenges underscore the complexity of adapting language model paradigms to the chemical domain and highlight the need for innovative approaches in molecular representation, data acquisition, and model evaluation.

### What’s special about FragLlama

The FragLlama model adapts the next-token prediction paradigm used in decoder-only Large Language Models (LLMs) to molecular design. In this approach, we introduced new tokens specifically trained to represent molecular fragments and functional groups. The current token vocabulary consists of 8,000 tokens, model trained on approximately 70 billion tokens, with the model size set at 779 million parameters. The model is designed to handle tasks such as fragment growth, scaffold modification, scaffold hopping, and linker design. By framing molecular design as a next-fragment prediction task, FragLlama mimics the strategies employed by medicinal chemists, who often grow molecules from a starting fragment or substitute functional groups on a core scaffold. We trained the tokens from scratch, ensuring they are tailored to better understand molecular fragments and functional groups, rather than using tokens from LLMs trained on human language or SMILES atom tokens.

Building upon this innovative framework, we explored FragLlama’s practical applications in diverse molecular design scenarios. The model demonstrates its ability to generate diverse and chemically valid structures, exploring both known and novel chemical spaces. We illustrate this through the creation of a molecular glue library, where FragLlama not only reproduces expert-designed compounds but also ventures into unexplored regions of chemical space. In PROTAC linker design, FragLlama exhibits proficiency in generating structurally diverse linkers, including rigid ones, which are particularly valuable in this field. Furthermore, we demonstrate FragLlama’s adaptability through fine-tuning experiments focused on EGFR inhibitor design. The model’s performance improves when provided with comprehensive input data, including known inhibitors and their bioassay results. Throughout these applications, FragLlama showcases its potential to assist medicinal chemists by generating expert-level designs while also offering novel structural ideas.

## Methods and Discussions

### Data preparation and tokenization to represent molecular fragments

Recent research has highlighted significant limitations in the SMILES (Simplified Molecular Input Line Entry System) ^20^ representation of molecules, particularly in the context of its application to language models. Several studies have identified a fundamental issue: the position of atoms in SMILES strings does not correspond to their spatial arrangement in chemical graph representations.^21–23^ This discrepancy introduces several potential problems:

1. Spatial coherence in token prediction: The next-token prediction paradigm fails to consider the correct spatial ”order” of molecular generation, potentially leading to chemically implausible structures.
2. Tokenization challenges: The lack of spatial correspondence hinders effective tokenization, which is crucial for language model performance.
3. Constraints on generation strategies: SMILES representation necessitates complete, global generation of the entire molecule, rather than allowing for fragment-based or incremental generation. This requirement significantly alters the embedding space and complexity of next-token prediction tasks. Furthermore, it imposes limitations on fragment insertion and other partial generation techniques.

To address these issues, we trained tokens from scratch to represent molecular fragments, intentionally incorporating chemical domain knowledge prior to the language model processing, effectively reducing the complexity of the next token prediction.

Additionally, we have implemented corresponding data augmentation algorithms. These algorithms ensure that high-quality datasets can be expanded while maintaining chemical validity. Our testing demonstrates that this approach allows for lossless reconstruction of the original SMILES molecule, preserving chemical integrity throughout the augmentation process.

We used the Byte Pair Encoding (BPE)^24^ algorithm for tokenization. Its fundamental principle can be described as follows: BPE initiates at the character level and proceeds through iterative merges of the most frequently occurring adjacent character pairs or sub-word pairs. Each iteration generates a new subword, which is subsequently added to the vocabulary. This process continues until a predefined vocabulary size is reached or a stopping criterion is met.

The efficacy of BPE lies in its ability to create a vocabulary that efficiently represents the data while balancing the trade-off between vocabulary size and token length. In the context of chemical representations, BPE can capture meaningful substructures and patterns within molecular strings, potentially leading to more effective encoding of chemical information for language models. This approach allows for adaptive tokenization that can reflect the underlying structure and frequency of molecular subunits, potentially enhancing the model’s ability to learn and generalize chemical patterns.

Figure 1 illustrates the process of our fragment-based BPE (Byte Pair Encoding) training. Unlike traditional methods, our approach incorporates an additional preprocessing step, which segments input molecules into smaller fragments before applying BPE. This allows the algorithm to recognize common patterns at a more granular level, leading to an optimized vocabulary specifically suited for molecular structures. The strength of this method lies in its ability to more effectively tokenize new and unseen molecules, enhancing the model’s generalization capabilities.

**Figure 1:**
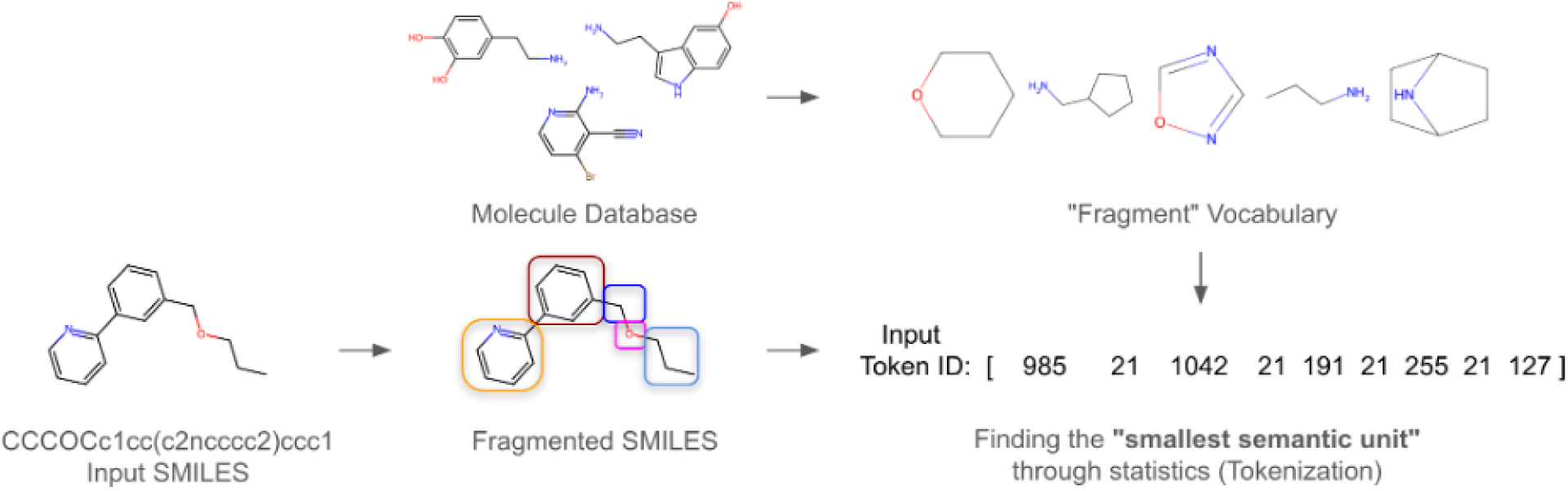
Fragment-level tokenization process for molecular design. Starting from input SMILES strings, molecules are retrieved from a molecular database and fragmented into smaller units. These fragments form a ”fragment vocabulary”. In the tokenization process, the input SMILES is broken down into these molecular fragments, each assigned a corresponding token ID. The process identifies the ”smallest semantic unit” for molecular representation through statistical methods, facilitating the generation and manipulation of molecular structures in subsequent tasks.

### FragLlama Model Design

#### Next Token Prediction for Molecule Language Modeling

Next Token Prediction is the core task of large language models (LLMs). In this task, the model predicts the most likely next word/token based on a given context sequence. By analyzing linguistic patterns and semantic relationships in existing text, the model learns to predict the subsequent elements in a sequence. This process is autoregressive, meaning the model generates tokens sequentially, incorporating each newly generated token into the context for subsequent predictions. Next Token Prediction enables LLMs to generate coherent text, complete sentences, and excel in various natural language processing tasks.

The transformation of complex human language and thought into Next Token Prediction represents a remarkably ingenious concept. The loss function employed is simply cross-entropy, a fundamental measure in information theory. Entropy and information are essentially manifestations of ”non-uniformity.” Cross-entropy can be viewed as a form of ”compression assistance.” In the context of Molecule Large Language Models, for example, the token following a double bond in SMILES notation (represented by an equals sign ”=”) is not arbitrary, but rather represents ”non-uniform” information. The entire model training process can be conceptualized as the model’s progression from a ”uniform” next token prediction to a non-uniform state. In LLMs, the metric ”perplexity” is used to characterize the confidence level of the next token prediction. We can readily observe that the decrease in loss value tends to plateau early, while internal learning within the model continues. This process reflects the model’s transition from a state of ”ignorance” to one of ”knowledge.” In its initial state, the model’s predictions for all tokens are uniform, representing maximum uncertainty and minimum information content. As training progresses, the model’s predictions become increasingly ”non-uniform,” signifying greater certainty and information richness.

In the chemistry domain, this translates to the model learning the inherent constraints and patterns of molecular structures. For instance, the model learns that certain atomic combinations or bond types are more probable in specific contexts, mirroring the non-uniform distribution of chemical structures in reality. This transformation not only enables the model to accurately predict the next token but also allows it to generate meaningful and chemically plausible molecular structures. This process reflects a fundamental aspect of machine learning in complex domains, where the transition from a state of maximum entropy to one of structured knowledge mirrors the acquisition of domain-specific understanding.

The elegance of this approach lies in its ability to capture complex chemical knowledge through a conceptually simple prediction task. By minimizing cross-entropy, the model is effectively learning to compress chemical information efficiently, capturing the underlying patterns and rules of molecular structures. This framework allows for the implicit learning of chemical principles without the need for explicit rule encoding, potentially enabling the model to discover novel patterns or relationships in chemical data.

Figure 2 illustrates our FragLlama model architecture, which is modified from the Llama 3 architecture,^25^ and demonstrates how the classic ”Next token prediction” task is applied to this fragment-based molecule language model.

**Figure 2:**
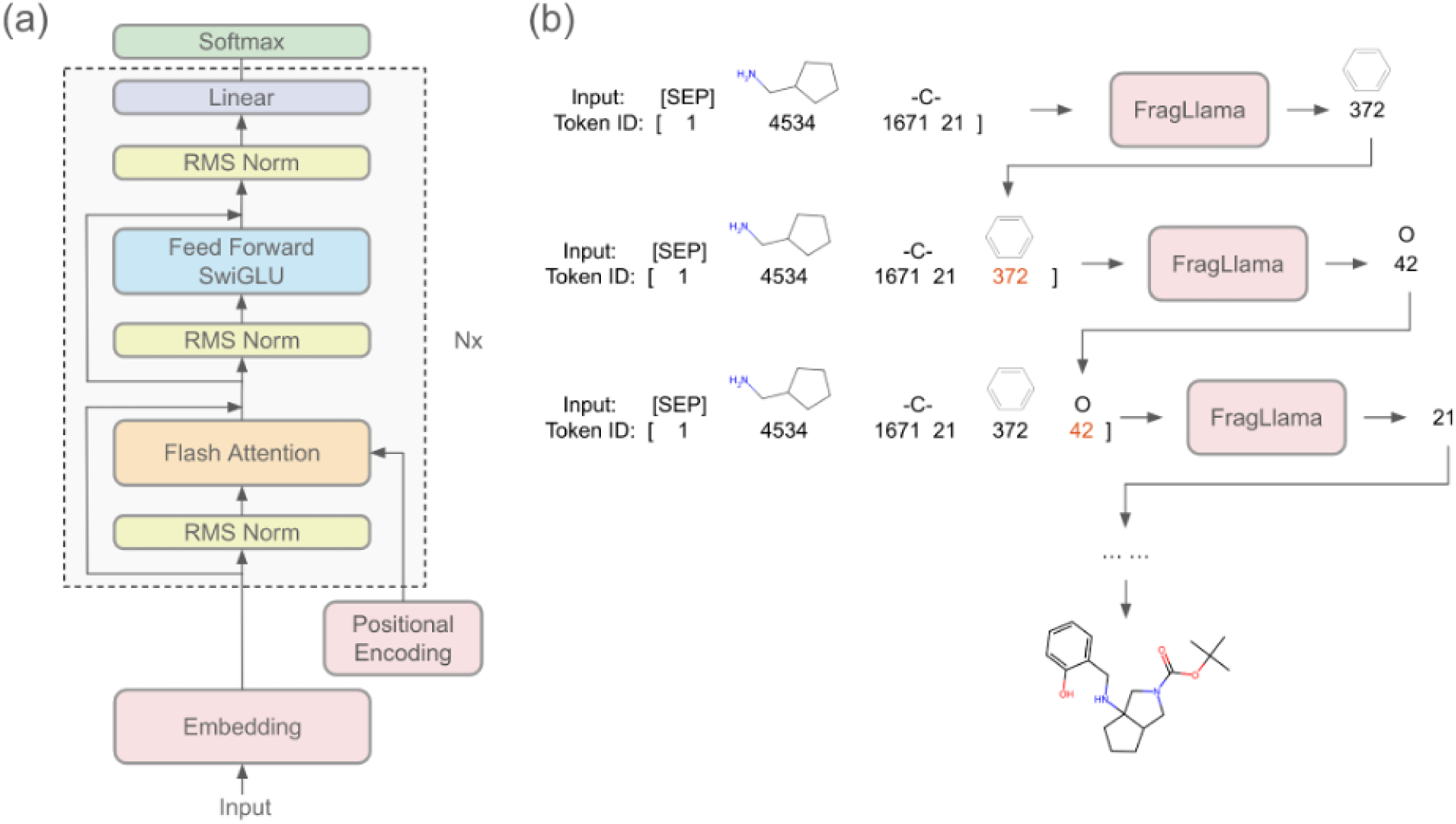
FragLlama architecture and molecular fragment generation process. (a) The architecture of the FragLlama model, built on a multi-layer transformer framework with components including Embedding, Positional Encoding, Flash Attention, SwiGLU Feed Forward layers, and RMS normalization. The architecture repeats Nx times, allowing for deeper processing of molecular fragment representations. (b) The fragment-level token generation process. Starting with an initial input molecule, FragLlama predicts the next fragment token iteratively. Each step adds a new fragment or functional group, progressively generating a complete molecular structure. Token IDs corresponding to molecular fragments are shown, and new tokens predicted at each step are highlighted in red.

#### The Transformer Architecture and Why We Choose Decoder-Only LM

The Transformer architecture, introduced by Vaswani et al. in 2017,^9^ represents a paradigm shift in neural network design for sequence processing. This architecture eschews traditional recurrent or convolutional structures in favor of a mechanism based entirely on attention to capture dependencies within input sequences. The core components of the Transformer include Multi-Head Attention (MHA) layers and Feed-Forward Network (FFN) layers. This design enables parallel processing of input sequences, significantly enhancing training efficiency and model performance. Transformers typically employ an encoder-decoder structure, though variations exist.

The Transformer’s powerful feature extraction capabilities and scalability have established it as the foundational architecture for modern natural language processing tasks and multimodal applications. In the biological and chemical domains, the most notable application is the AlphaFold series,^26,27^ which has revolutionized protein structure and biomolecular interaction prediction.

While the original Transformer architecture introduced in the seminal paper was an encoder-decoder structure, subsequent developments have led to the widespread adoption of encoder-only and decoder-only architectures:

1. Encoder-Only Models: These are primarily used for global sequence embedding. In biology, examples include the ESM series of protein language models. ^28^ In NLP, the classic example is Sentence BERT. These models excel at capturing contextual representations of entire sequences.
2. Decoder-Only Models: These are more suited for generalized few-shot learning tasks. Through techniques like prompting, they can be adapted to various domains without fine-tuning. The most prominent example is the GPT series. These models are particularly adept at generative tasks and can be more flexible in their applications. Decoder-only Models can be treated as Encoder-Decoder Models but share the same weights across Encoder and Decoder.

Generally, at smaller model sizes, BERT-like encoder-only models tend to outperform GPT-like decoder-only models, as the autoregressive loss function in decoder-only models is more challenging to optimize. ^29^ However, as model size increases, decoder-only models often demonstrate superior performance and versatility. Our choice of a decoder-only model architecture for molecule language modeling is motivated by its greater flexibility and the relative novelty of this approach in the field. To the best of our knowledge, many aspects of chemistry language modeling remain unexplored. This choice allows us to leverage the generative capabilities and adaptability of decoder-only models in tackling various chemistry-related tasks, from molecular property prediction to drug design.

#### Main Architectural Differences between FragLlama and Llama 3

The Llama series of large language models, developed by Meta AI (formerly Facebook AI Research), represents a significant advancement in the field of natural language processing. The Llama architecture is based on the decoder portion of the Transformer. Compared to GPT2, Llama incorporates several crucial improvements and optimizations:^25^

1. Rotary Position Encoding (RoPE):^30^ This technique efficiently handles positional information in the input sequence, allowing the model to better understand the relative positions of tokens.
2. SwiGLU Activation Function:^31^ This enhanced activation function improves the model’s capacity for non-linear transformations, potentially leading to better performance across various tasks.
3. Grouped Query Attention (GQA):^32^ Optimizations in the attention mechanism contribute to the model’s computational efficiency by reducing the KV cache.

These architectural choices enable Llama to demonstrate robust performance across a wide range of natural language processing tasks while maintaining relatively modest model sizes and computational requirements.

The reason we chose Llama is that, unlike some closed-source models, Llama’s openness promotes open research and innovation in the AI community. Although we did not utilize Llama weights directly, our model development was significantly informed by research that employed Llama weights, such as Alpaca,^33^ TinyLlama,^34^ and QLoRA.^35^ These studies provided substantial insights and inspiration for our approach to model development. The advancements made in these Llama-based models offered valuable lessons in efficient training techniques, model architecture optimization, and fine-tuning strategies. By leveraging the Llama architecture for our molecule language model, we aim to combine the power of state-of-the-art natural language processing with the specific requirements of molecular representation and prediction. This approach allows us to benefit from the extensive research and optimization that has gone into Llama’s development while tailoring the model to the unique challenges of molecule language modeling.

Based on the specific requirements of small molecule generation and the enhanced efficiency of our tokenizer, we have implemented several modifications to the Llama 3 architecture. These adjustments are primarily driven by the following considerations:

1. Positional Encoding: We have replaced the Rotary Position Encoding (RoPE) with a standard positional encoding scheme. This change is justified by the fact that our objective focuses on small molecule generation, where token sequences rarely exceed 100 tokens. The long-range context capabilities provided by RoPE, while beneficial for general language tasks, are not critical for our application. This modification simplifies our model architecture without compromising performance in our specific domain.
2. Attention Mechanism: We have opted to use Flash Attention instead of the Grouped Query Attention (GQA) employed in Llama 3. While GQA offers computational efficiency advantages and facilitates single-card inference for larger models, it can potentially reduce model performance. The trade-off introduced by GQA is more beneficial for models with 7 billion parameters or more, where the reduction in attention parameters can be compensated by increased Feed-Forward Network (FFN) capacity. However, for our relatively smaller model size, this trade-off is suboptimal.

Flash Attention^36^ provides an efficient attention computation without the performance drawbacks of GQA. This choice allows us to maintain the full expressiveness of the attention mechanism, which is crucial for capturing the intricate patterns and relationships in molecular structures.

By carefully considering the trade-offs between model complexity, performance, and computational efficiency in the context of molecular generation, we have developed a more targeted and potentially more effective architecture for molecule language models. This tailored architectural modification simplifies the model structure and improves training efficiency. The use of Flash Attention ensures that we do not sacrifice model quality for computational efficiency, which is particularly important given our smaller model size. These changes reflect careful consideration of the unique characteristics of molecular data and the specific requirements of molecule language models.

### Pre-Training

Our training infrastructure, supported by the Oracle Cloud Team, represents a significant computational resource tailored for the demands of large-scale language model training in the domain of chemistry. We utilized a cluster of 7 NVIDIA A100 GPUs for the pre-training phase. And using DeepSpeed with ZeRO Stage 3 optimization for distributed training.

### Alignment–Fine Tuning

Alignment in LLM refers to the crucial challenge of ensuring AI systems behave in ways that are consistent with human values and intentions, especially as these systems become more advanced and autonomous. Researchers and developers employ various methods to address this challenge, including fine-tuning pre-trained models on specific datasets, Reinforcement Learning from Human Feedback (RLHF)^37^ to guide models using human preferences, reward modeling to accurately reflect human values.

Alignment in LLMs also equips molecular LLMs with the ability to incorporate various types of extra information, such as experimental data, modeling data, and prior data, into the molecule generation process. By integrating this additional data, molecular LLMs can align more closely with real-world constraints and objectives, enhancing their predictive accuracy and relevance in drug discovery. For instance, experimental data on compound potency or selectivity can be used to fine-tune the model, ensuring that generated molecules meet specific biological criteria. Similarly, modeling data, such as structure-activity relationships (SAR) or docking scores, can guide the generation toward favorable chemical spaces. Prior data, representing previously collected datasets or knowledge, helps further refine the model’s output, ensuring it builds upon existing research to generate novel, high-potential compounds. This ability to integrate diverse sources of information is a key advantage of alignment in molecular LLMs, making them more adaptive and useful for real-world drug design challenges.

To demonstrate our current model’s significant fine-tuning potential, we conducted a proof-of-concept experiment. Our objective was to investigate whether the fine-tuned model could generate molecules more closely resembling human-designed EGFR binders. Here, we collected 8,532 EGFR inhibitors from PubChem with active bioassay results and valid IC50 values to fine-tune FragLlama. The dataset included approximately 6,822 molecules with high IC50 values and 1,710 molecules with low IC50 values. Two distinct fine-tuning approaches were implemented:

1. The first approach involved direct utilization of the 6,822 molecules with high IC50 values.
2. The second approach combined all molecules, incorporating an untrained special token at the beginning of each molecule to denote its IC50 range.

Given the nature of fine-tuning, we employed a reduced learning rate and a shorter training duration. This strategy was aimed at preserving the model’s pre-trained knowledge while adapting it to the specific EGFR binder-related chemical space. For molecular design, we selected to use the scaffold 4-Anilinoquinazoline as its the most common smallest scaffold in our curated dataset. Given the multiple growth anchors identified in this scaffold, we didn’t assign specific growth anchors. Accordingly, we conducted three types of molecular design: 1) default settings without fine-tuning; 2) fine-tuned with 6,822 EGFR inhibitors, without IC50 special tokens; 3) fine-tuned with both 8,532 EGFR inhibitors and their IC50 special tokens.

#### Decoding Strategies

LLM decoding strategies are methods for controlling the quality and diversity of output when generating text from language models. These strategies determine how to select the next token from the model’s probability distribution. The specific decoding strategy we used is beam search variance, as it maintains multiple candidate sequences and ultimately selects the sequence with the highest overall probability. Common decoding strategies include Greedy Search, Beam Search,^38^ Temperature Sampling,^39^ Top-k sampling,^40^ Top-p (nucleus sampling),^41^ frequency penalty,^42^ and presence penalty.^10^ These strategies balance determinism and creativity in text generation to varying degrees and are suitable for different application scenarios. Choosing an appropriate decoding strategy is crucial for generating high-quality, coherent, and diverse text.

## Results and Discussion

### Tokenizer Compression Rate and Meaningful Tokens

In our study, we explored the efficiency of various tokenizers in representing chemical structures by evaluating their compression rates. The compression rate was defined as the ratio of the original molecular data size to the tokenized data size. A lower compression rate indicates that the tokenizer is more efficient in representing the molecular structures with fewer tokens, while a higher compression rate suggests the opposite.

Take the molecule illustrated in Figure 3, a GLP-1R inhibitor^43^ developed by Gilead Science, as an example. Using the FragLlama fragment-level tokenizer with an 8k vocab size, the resulting token length is 61. The token sequence length obtained using the atom-level tokenizer is 105 (the atom-level tokenizer is simply char-level tokens with regex match). The GPT2 tokenizer with a 5k vocab size yields a token length of 86, while the GPT4 tokenizer with a 128k vocab size results in a length of 71.

**Figure 3:**
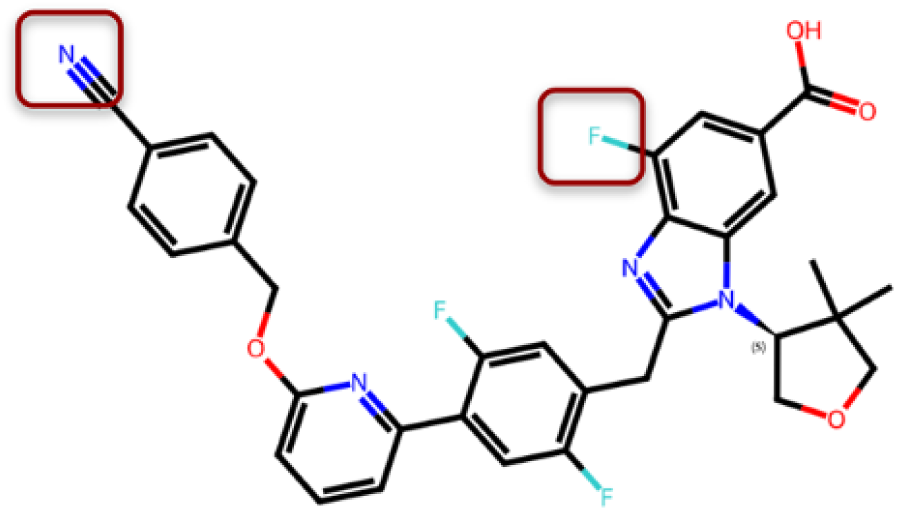
A GLP-1R inhibitor (Gilead Science)

From the current perspective, we have achieved a better ”compression ratio” than GPT4 using an 8k vocab size. Considering only the compression ratio, shorter sequences bring many benefits. For instance, for molecules similar to the GLP-1R inhibitor, the FragLlama tokenizer needs to perform 61 next token predictions, while the atom-level tokenizer requires 105. Additionally, shorter input sequences for molecules of the same length mean less memory usage and more efficient attention queries. This allows models with the same architecture and number of parameters to easily handle more complex input sequences.

Secondly, for a tokenizer, the ”semantic representation ability” is more important compared to the compression ratio. Although GPT4’s token compression capability for GLP-1R inhibitor is similar to the FragLlama Tokenizer, GPT4’s training data is mainly based on human language, which means GPT4’s tokens lack chemical significance. Based on prior chemical knowledge and statistical findings, our ”molecular language tokens” can be visual-ized, and they are all common components of drug molecules. This is not surprising, as the ”natural language tokens” obtained by LLMs like Llama through statistical patterns also usually have linguistic value. Excellent semantic representation ability is one of the core factors in making the model easy to train and reducing model complexity. In comparison, although the atom token has a smaller vocab size, the diversity of SMILES makes the model much more difficult to train than the FragLlama Tokenizer. For example, to generate a benzene ring, the atom-level tokenizer requires 6 times more memory in the attention layers, and needs to perform 6 times of next token predictions, and this is assuming the optimal decoding parameters have been found, which adds a lot of potential instability.

Figure 4 showcases some typical tokens used by the FragLlama model. Although FragLlama’s Tokenizer is built based on the BPE (Byte Pair Encoding) algorithm, after training, we can observe that many of the generated tokens actually correspond directly to fragments of chemical molecules. This phenomenon bears an interesting similarity to the results of BPE training in human language processing. Just as in natural language processing, where tokens after BPE training often retain linguistically meaningful units (such as roots, affixes), FragLlama’s Tokenizer has successfully captured meaningful fragments in chemical structures. This result suggests that our method is not merely a mechanical segmentation of SMILES strings, but has successfully learned the chemical semantics within molecular structures. This chemically informed segmentation approach helps the model better understand and generate molecular structures, as it directly operates on chemically meaningful units rather than just abstract character sequences.

**Figure 4:**
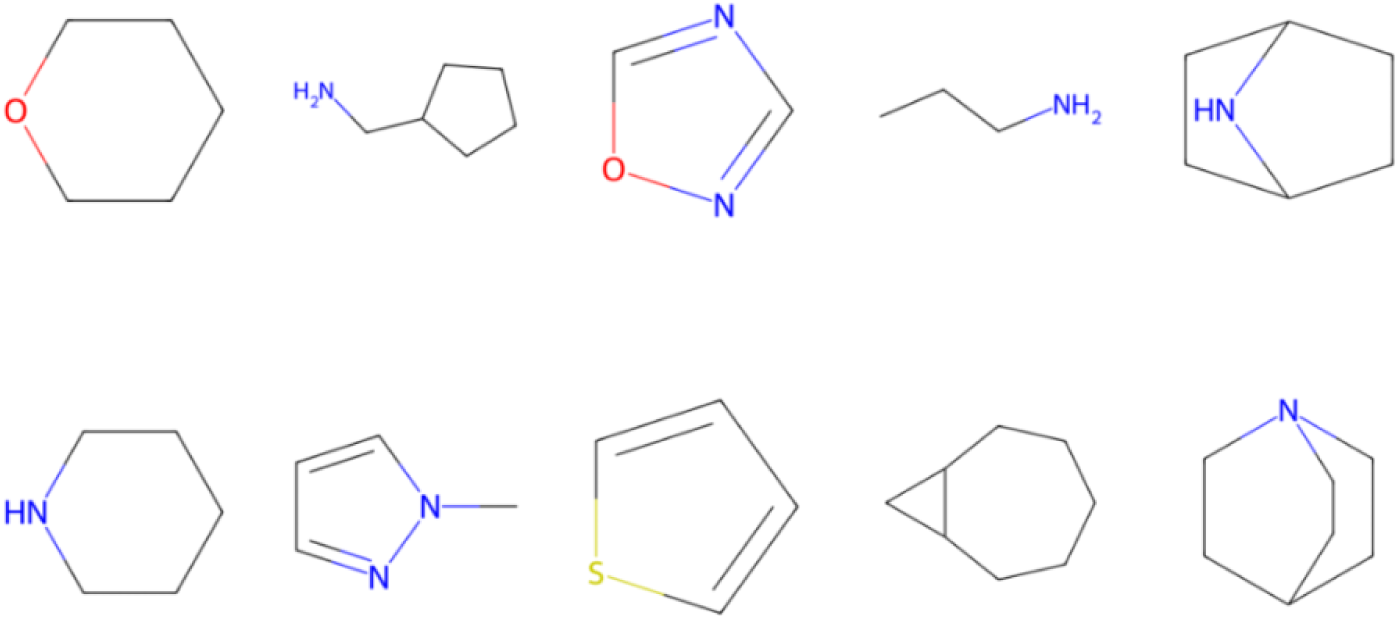
FragLlama Token Visualization

We have repeatedly witnessed the instability of prompts in LLMs like GPT, which is because human language itself is highly variable, and a change in a single word can completely alter the expressed meaning. We believe this is also true for molecular language (SMILES). A better token vocab set can effectively alleviate this issue.

In summary, a good tokenizer significantly reduces vocabulary size while maintaining good semantic representation ability, which helps to reduce model complexity and memory requirements. Secondly, it can better handle unknown and rare words (in our cases, the uncommon molecules), improving the model’s generalization ability. Moreover, an efficient tokenizer can speed up model training and inference. A good tokenizer can also preserve important linguistic information through reasonable segmentation methods. In the specific domain of molecular design, our tokenizer is more conducive to preserving chemical functional groups and capturing atomic connectivity patterns. This ”linguistic” parsing approach to chemical structures enables the model to directly manipulate chemically meaningful units, rather than just abstract character sequences, thereby improving the model’s performance and efficiency in molecular design tasks and laying a solid foundation for subsequent model designs.

#### Attention Weights Visualization

The attention mechanism allows the model to focus on different parts of the input sequence when generating a particular output. It computes a weighted sum of the input tokens based on their relevance to each other, allowing the model to capture dependencies between tokens regardless of their distance from one another. Multihead attention was used to capture multiple types of relationships or features across the input sequence. Each head in multihead attention learns a different perspective on the input. By having multiple heads, the model can look at various aspects of the data simultaneously. It can learn both fine-grained and high-level patterns across different parts of the input sequence. After each head processes the input, the results are concatenated and combined into a single output, providing a rich, multi-faceted representation of the input sequence.

For instance, in the sentence, ”The scientist carefully analyzed the results of the experiment to ensure the accuracy of the findings, despite encountering several unexpected challenges along the way,” one head might focus on the grammatical structure, identifying subject-verb-object relationships such as the link between ”scientist,” ”analyzed,” and ”results.” Another head might track long-range dependencies, connecting the main clause with the subordinate clause by linking ”analyzed” with ”to ensure” to understand how the action relates to ensuring accuracy. Other heads may capture semantic roles, identifying the ”scientist” as the agent and the ”results” as the theme, or modifiers like ”carefully,” which gives more detail on how the action was performed. Additional heads focus on prepositional phrases like ”of the experiment” to capture spatial and causal relationships, sentiment and connotation to identify the contrast introduced by ”despite encountering several unexpected challenges,” and temporal or sequential relationships to follow the logical flow of events from analyzing the results to ensuring their accuracy. Some heads handle coreference resolution, linking ”results” to ”findings,” while others focus on contrast and concession, capturing the opposition between ensuring accuracy and encountering challenges.

In analogy to molecules, different attention heads in the FragLlama model may specialize in tracking various molecular features. For instance, one head might focus on the structural relationships between molecular fragments, identifying how different atoms and bonds contribute to synthesizability. Another head could track long-range dependencies between functional groups to predict lipophilicity, ensuring that hydrophobic and hydrophilic regions are properly represented. Similarly, other heads may capture chemical reactivity, such as tracking electrophilicity to determine how reactive certain parts of the molecule are. Just as in language models, each attention head plays a specialized role in interpreting different aspects of the molecular structure.

While models like GPT are highly powerful for linguistic tasks, the tokens and attention mechanisms they learn are not well-suited for tasks like molecular design and property prediction. The token embedding space in natural language models is optimized for capturing relationships between words, phrases, and syntax, which differ significantly from the structural and chemical properties of molecules. In molecular design, precise attention to interactions, bonding patterns, and chemical properties is critical. The role of attention in molecular models like FragLlama is to rearrange and combine molecular tokens in a way that captures these complex relationships, producing more meaningful representations for molecules. As a result, models trained on language data, like GPT2, are not interchangeable with those trained on molecular data, as they do not adequately capture the nuances of molecular features necessary for accurate predictions.

To explore this hypothesis, we used the GLP-1R inhibitor (Figure 3), comparing attention patterns between FragLlama and GPT2. We selected two tokens present in both the FragLlama Tokenizer and the GPT2 Tokenizer. As demonstrated in Figure 5, FragLlama employs a more targeted and efficient attention mechanism compared to GPT2. For token ’F’, FragLlama exhibits a highly selective attention pattern, concentrating its focus on critical heads, as evidenced by the sparse yet focused activations in panel (a). This selective attention likely reflects a refined mechanism tailored for molecular design tasks, where precision in capturing key features and relations among the tokens is paramount. In contrast, GPT2 displays a more diffuse attention pattern for the same token, which may be less optimal for molecular applications where attention needs to be sharply focused on specific molecular features. For token ’N’, FragLlama again demonstrates its ability to allocate attention selectively and effectively across fewer heads, as seen in panel (b). This contrasts with GPT2’s broader and more distributed attention in panels (c) and (d), which might dilute the model’s ability to focus on the most relevant aspects of molecular structures. The targeted attention strategy of FragLlama suggests that it is better suited for handling the specialized demands of molecular design.

**Figure 5:**
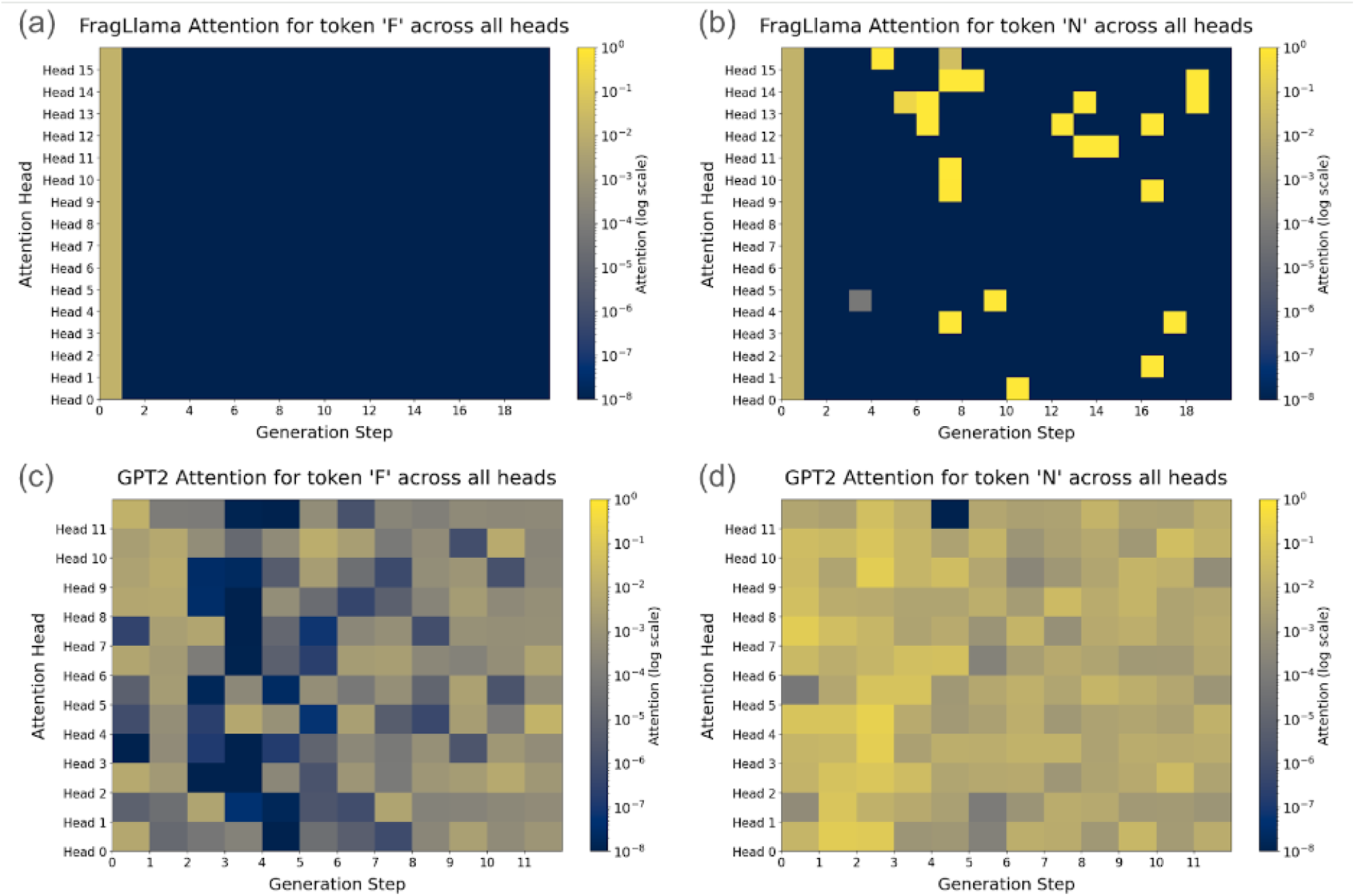
Attention patterns of FragLlama and GPT2 across heads for tokens ’F’ and ’N’. (a) FragLlama attention for token ’F’ across all heads and generation steps. (b) FragLlama attention for token ’N’ across all heads and generation steps. (c) GPT2 attention for token ’F’ across all heads and generation steps. (d) GPT2 attention for token ’N’ across all heads and generation steps. The color scale represents attention intensity on a logarithmic scale, with yellow indicating higher attention and blue indicating lower attention.

#### Fragment Growth Molecular Design: Showcasing the Creation of a Molecular Glue Library

Molecular glue degraders are small molecules that facilitate the targeted degradation of specific proteins by promoting the formation of a ternary complex between a target protein and an E3 ubiquitin ligase.^44,45^ Unlike traditional inhibitors that block the function of a protein, molecular glue degraders act as catalytic agents, enhancing the recruitment of an E3 ligase to the target protein, which is then ubiquitinated and directed to the proteasome for degradation. These degraders represent a novel therapeutic strategy in drug discovery, particularly valuable in targeting proteins that were previously considered ”undruggable”.^46^ Molecular glues have gained significant attention due to their potential to modulate protein interactions in a highly specific manner. For example, thalidomide ^47^ analogs like lenalidomide ^48^ and pomalidomide ^49^ are well-known cereblon binders, acting as molecular glues by bridging interactions between cereblon (CRBN) and transcription factors like Ikaros, leading to their ubiquitination and degradation. This mechanism not only opens new pathways for drug discovery but also enables the modulation of biological processes that are otherwise difficult to influence through traditional small molecule inhibitors. ^50,51^

We employed the fragment growth strategy provided by FragLlama to design molecular glues, using three common cereblon binders—S-Pomalidomide, S-Lenalidomide, and S-Thalidomide—as starting points. We identified 3, 3, and 4 fragment growth anchors in the Protein Data Bank (PDB) for S-Pomalidomide, S-Lenalidomide, and S-Thalidomide, respectively (Figure 6). Using our pretrained model for molecular growth from given fragments and anchors, we generated 5,834 molecules after sanitization and deduplication. To compare the distribution of these FragLlama-generated molecules in chemical space with expert designs, we curated a reference dataset of 73,443 molecules from PubChem containing substructures of these cereblon binders.

**Figure 6:**
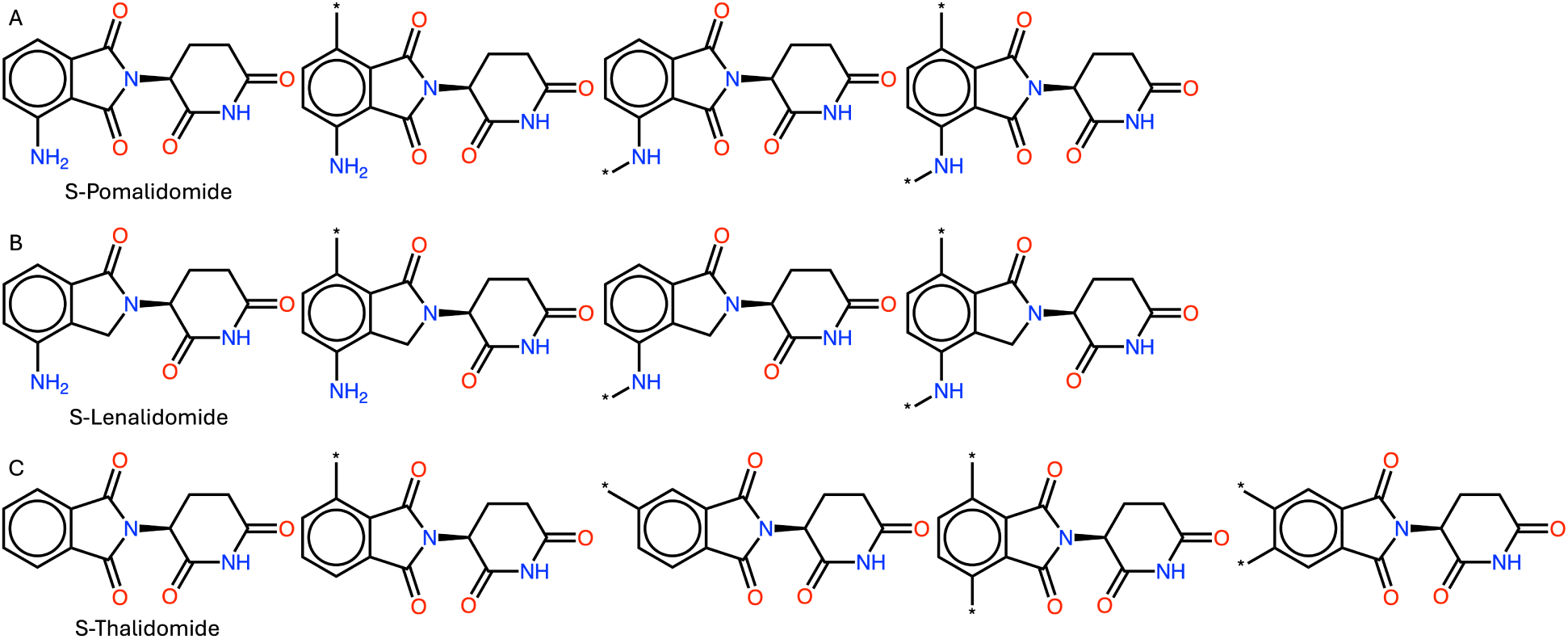
Fragment growth anchors for three cereblon binders: (A) S-Pomalidomide, (B) S-Lenalidomide, and (C) S-Thalidomide. The molecular structures of each cereblon binder are shown with their respective fragment growth anchors highlighted. These anchors serve as starting points for fragment growth.

We calculated Atom-pair descriptors ^52^ for both the curated PubChem molecules and FragLlama-designed molecules. Tanimoto distance^53^ was used to calculate pairwise similarities between fingerprints. To visualize and compare the distributions in chemical space, we applied UMAP (Uniform Manifold Approximation and Projection)^54^ for dimension reduction, creating a two-dimensional embedding.

The UMAP visualization (Figure 7) reveals that FragLlama-designed molecules partially overlap with expert-designed compounds from PubChem in chemical space. While PubChem compounds cover a broader range (PROTACs were included in the curated dataset), FragLlama also explores regions not represented in the curated dataset. Our analysis identified four distinct regions of novel chemical space that FragLlama successfully explored, with representative structures listed in Table 1. This demonstrates the model’s capability to both reproduce the distribution of known drug-like compounds and design innovative molecules in underexplored areas.

**Figure 7:**
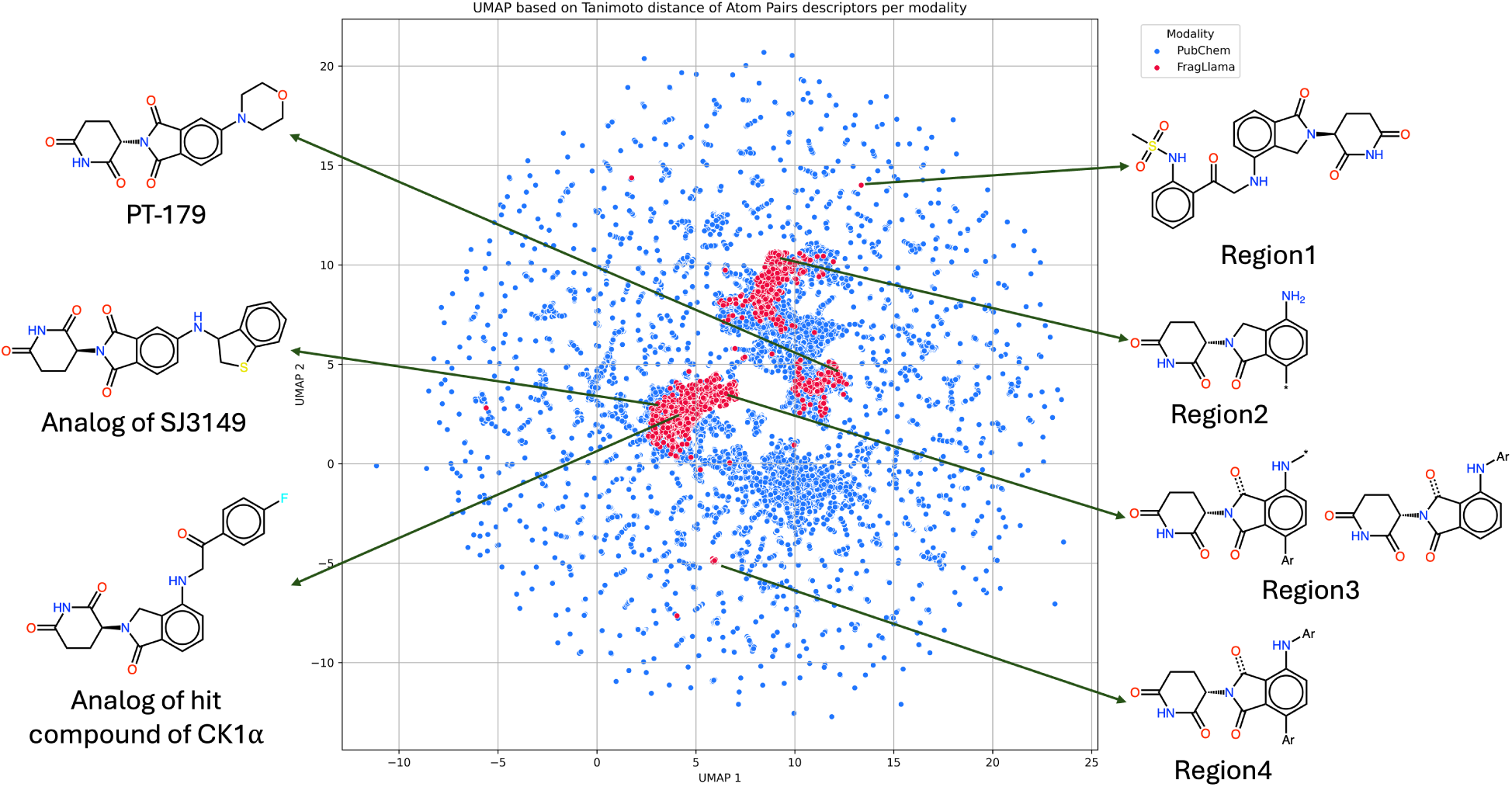
Visualization of molecular distribution in chemical space using Uniform Manifold Approximation and Projection (UMAP). The plot compares FragLlama-designed molecular glue molecules (red) with expert designed molecules, curated from PubChem (blue). Four novel chemical space regions (Regions 1-4) explored by FragLlama are highlighted on the right, with representative molecular structures illustrating key structural features. Three recently reported molecular glues are also indicated on the left: an analog of the hit compound for CK1*α* degradation, an analog of SJ3149 (optimized CK1*α* degrader), and PT-179 (a novel IMiD derivative)

**Table 1:**
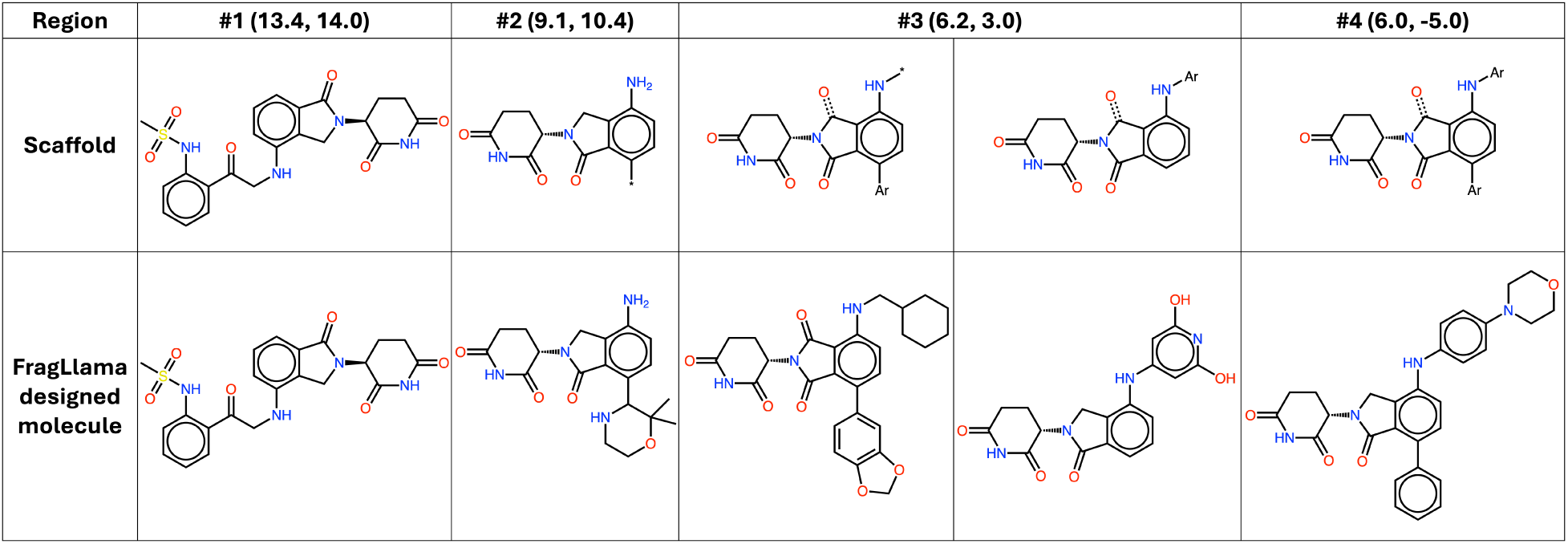
Comparison of scaffold structures and FragLlama-designed molecules across three distinct regions of chemical space. The top row shows the original scaffolds used as starting points in each region, while the bottom row presents the novel molecules generated by FragLlama in the corresponding regions. The regions, defined by specific coordinates, demonstrate FragLlama’s ability to explore new areas of chemical space and design structurally diverse molecules from the initial scaffolds.

To further validate FragLlama’s performance in designing drug-like molecules, we compared our generated compounds to recently reported molecular glues. Notably, we observed striking similarities between FragLlama’s designs and several recently disclosed molecular glues. For instance, a recent study described potent, selective, and orally bioavailable molecular glue degraders of CK1*α*, including a hit compound and its optimized version, SJ3149 (Figure 8A-B, first row).^55^ Remarkably, we identified a FragLlama-designed molecule ((4.2, 2.5) in Figure 7) that is structurally similar to the hit compound, with only one nitrogen atom moved one position along the chain of the molecule (Figure 8A, second row). Moreover, we found a FragLlama-designed molecule ((3.1, 2.9) in Figure 7) that is highly similar to SJ3149. The attached heterocyclic rings in both molecules follow a very similar pattern, with both featuring a five-membered ring fused to a six-membered aromatic ring and connectivity remain consistent, preserving the linkage through the NH group. This further demonstrates our model’s capacity to generate pharmaceutically relevant compounds. In another example, we compared FragLlama’s output to PT-179, a novel IMiD derivative that binds CRBN without inducing degradation of off-target proteins (Figure 8C, top).^56^ Once again, we identified a FragLlama-designed molecule ((12.2, 4.4) in Figure 7) identical to PT-179. These findings highlight FragLlama’s ability to design expert-level molecules, underscoring its value in real-world drug discovery efforts.

**Figure 8:**
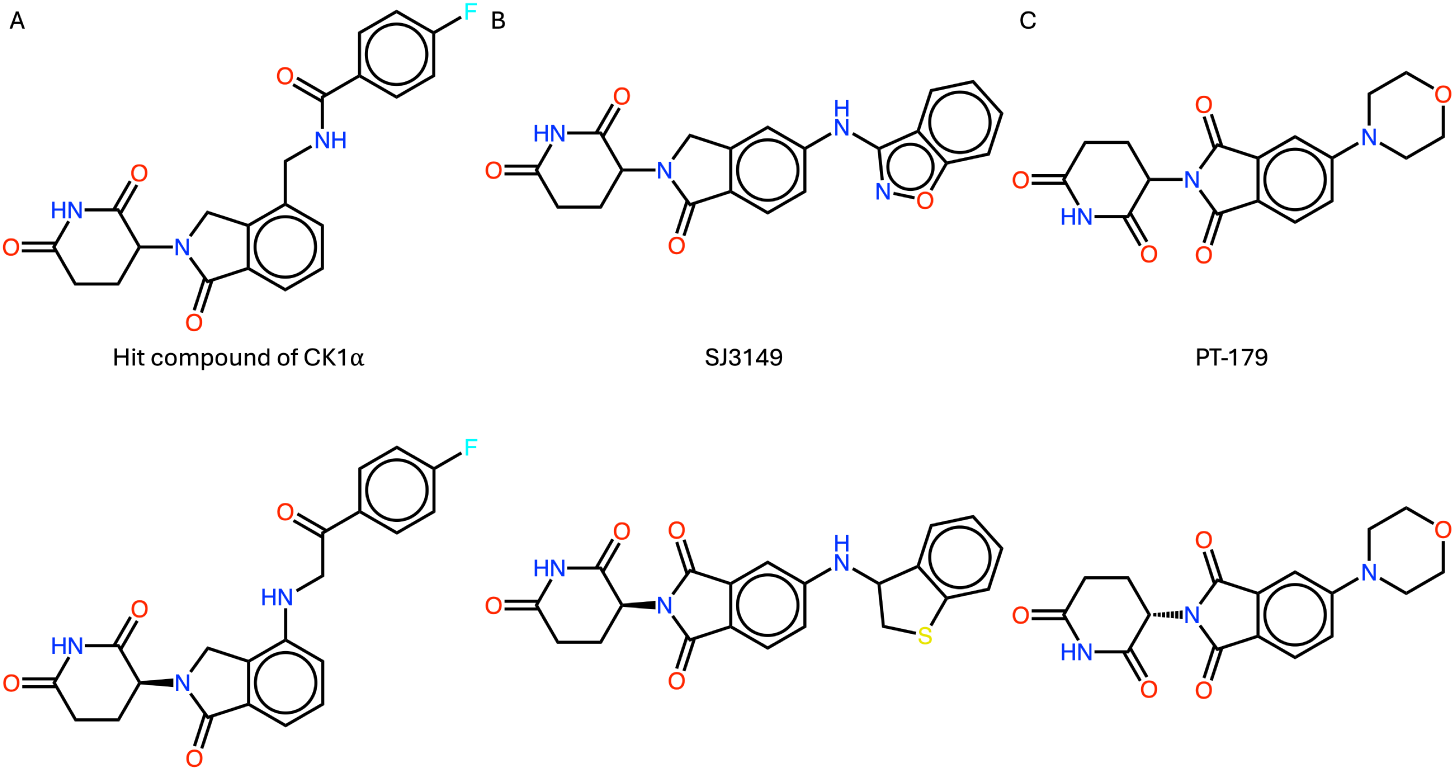
Comparison of recently reported molecular glues with FragLlama-designed molecules: (A) The hit compound for CK1*α* degradation and its structurally similar FragLlama-designed molecule. (B) SJ3149, an optimized CK1*α* degrader from the hit compound, alongside a structurally similar FragLlama-designed molecule. (C) PT-179, a novel IMiD derivative that binds CRBN without off-target protein degradation, and its structurally identical FragLlama designed molecule.

#### Fragment Linking Molecular Design: Showcasing PROTAC Linker Designs

PROTACs are another type of degrader that differ from molecular glue degraders in their mechanism of action. While molecular glues promote the natural interaction between a target protein and an E3 ubiquitin ligase, PROTACs are bifunctional molecules that chemically link the two together. One end of a PROTAC binds to the target protein, and the other binds to the E3 ligase, facilitating ubiquitination and subsequent degradation. A key aspect of PROTAC design is the linker that connects these two ligands; linker engineering is crucial for optimizing the spatial arrangement and stability of the ternary complex, directly impacting the efficiency of the degradation process.^57–60^

Here, we tested three PROTACs: SP27^61^ (inducing ternary structure with cereblon and PLK4 protein, Figure 9A), ARV-471^62^ (cereblon and estrogen receptor, Figure 9B), and CP07^63^ (cereblon and CDK9 protein, Figure 9C).

**Figure 9:**
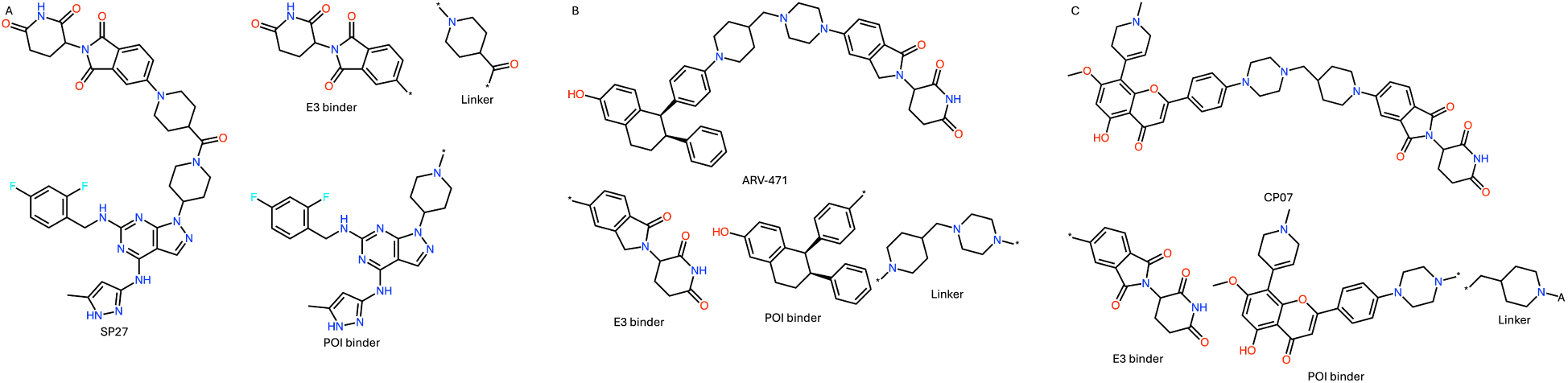
Structural representations of three PROTAC examples with their fragment growth anchors: (A) SP27 (cereblon-PLK4), (B) ARV-471 (cereblon-estrogen receptor), and (C) CP07 (cereblon-CDK9). The molecular structures highlight the E3 ligase binders, target protein binders, and linkers, with fragment growth anchors indicated for linker design by FragLlama.

FragLlama designed 156, 537, and 52 linkers for these PROTACs, respectively. To compare our designed linkers with the expert designed ones, we calculated similarities using Atom-pair descriptors^52^ for the whole molecular structures and Tanimoto distance ^53^ as the similarity metric. For SP27, we found a designed PROTAC with a similarity score of 0.93, reproducing both the carbonyl group and six-member ring, with the only structural difference being a missing nitrogen atom in the ring (Figure 10A). For ARV-471 and CP07, the most similarly designed PROTACs had scores of 0.78 and 0.74, respectively. While we didn’t reproduce the six-member rings, we achieved the same carbon chain lengths and generated branch chains with methyl and carbonyl groups to mimic the ring structure. (Figure 10B-C). These examples demonstrate FragLlama’s capability to design drug-like linkers for PROTACs. The distribution of similarity scores for these three examples showed a wide range, indicating high diversity in our designed linkers (Figure 11). This suggests that FragLlama can explore a wide range of linker structures in chemical space, potentially leading to the design of novel, effective PROTAC linkers. Rigid linkers are particularly favored in linker engineering. We also examined its capability of generating rigid linkers, as illustrated in Figure 12.

**Figure 10:**
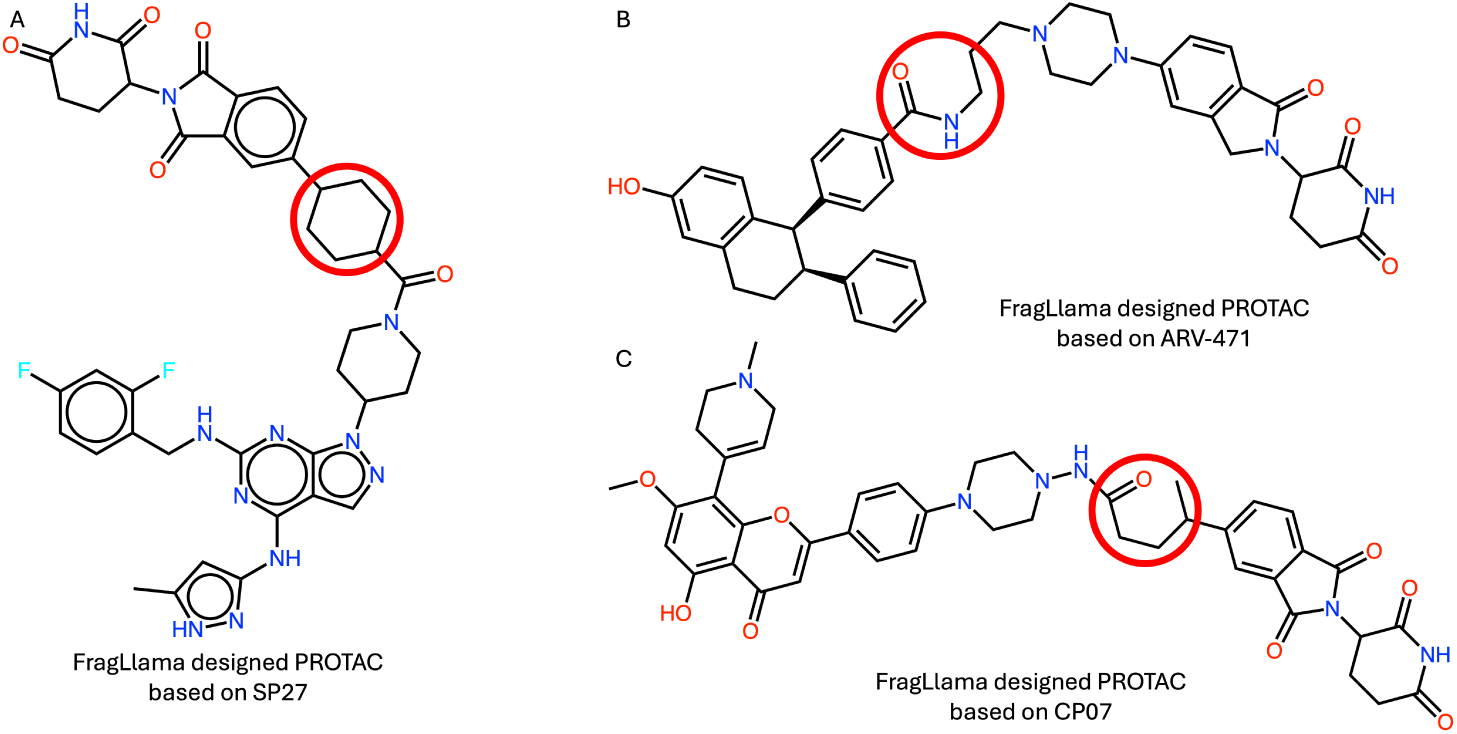
FragLlama-designed PROTACs with high structural similarity to reference PRO-TACs: (A) SP27 (similarity score: 0.93), (B) ARV-471 (similarity score: 0.78), and (C) CP07 (similarity score: 0.74).

**Figure 11:**
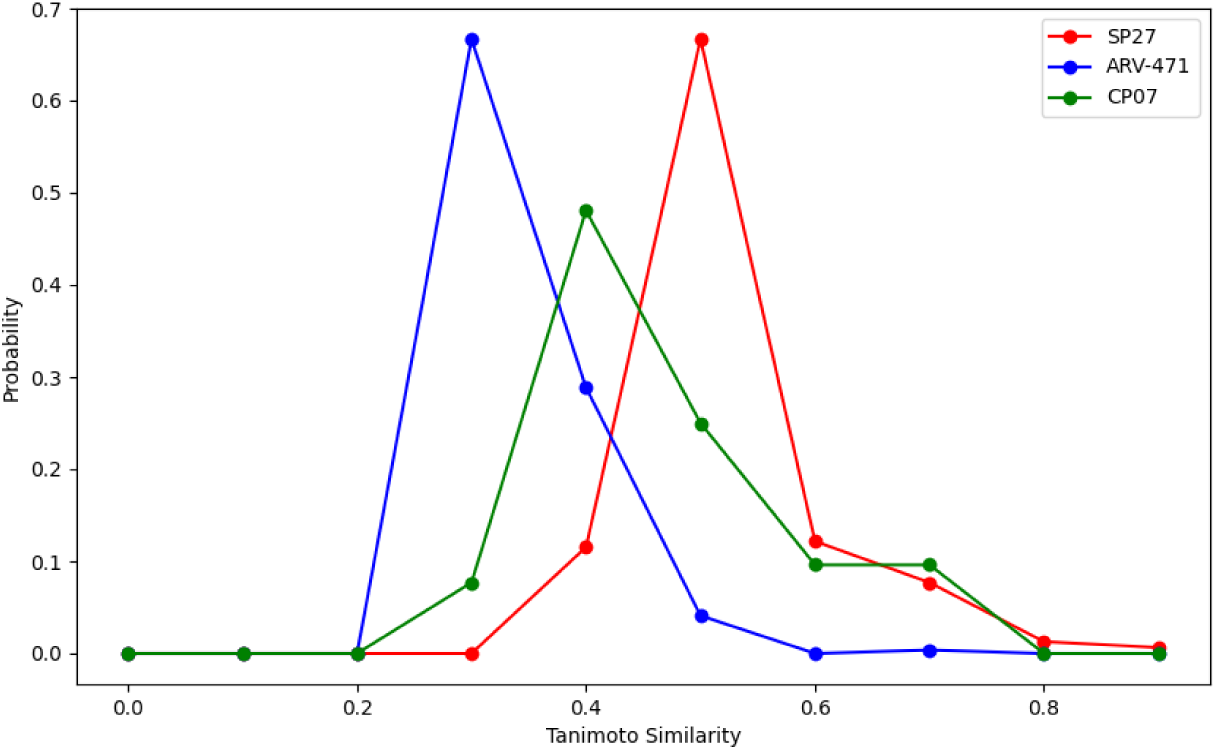
Distribution of Tanimoto similarity^53^ scores for FragLlama-designed PROTACs compared to three reference PROTACs: SP27 (red), ARV-471 (blue), and CP07 (green).

**Figure 12:**
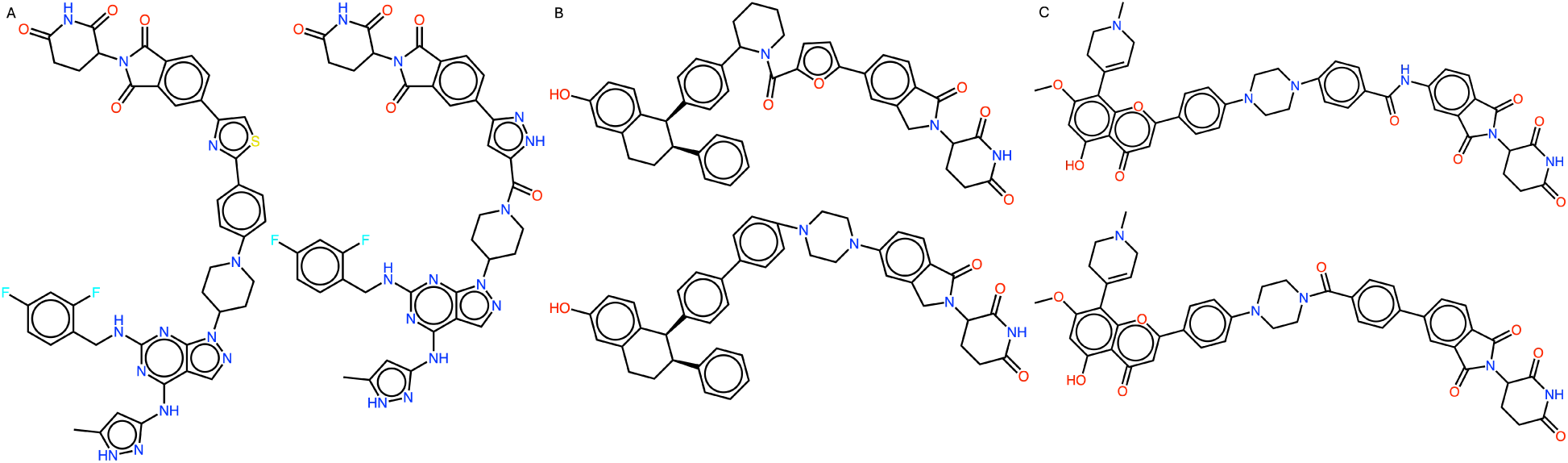
Examples of FragLlama-designed PROTACs incorporating rigid linkers, based on (A) SP27, (B) ARV-471, and (C) CP07 scaffolds.

#### Modify Molecules Towards Desired Properties: EGFR binder design

To compare FragLlama-designed molecules with known inhibitors, we selected 1,769 EGFR inhibitors from PubChem that have 4-Anilinoquinazoline. We visualized the results using UMAP (Figure 13). The UMAP visualization revealed that while all three sets of FragLlama-designed molecules occupied different chemical spaces than the PubChem compounds, the set designed with the model fine-tuned with EGFR binders and IC50 tokens showed greater overlap with PubChem regions. The binder-fine-tuned-only set performed better than the non-fine-tuned set in this regard. The challenge for FragLlama lay in designing drug-like compounds with greater complexity, as the scaffold we used was the smallest common structure, while real compounds in the PubChem dataset were more elaborate. This explains why FragLlama-designed molecules without fine-tuning remained largely separate from PubChem-covered regions. However, with a fine-tuned model with both binders and IC50 tokens, FragLlama successfully designed some molecules within PubChem-covered regions. Table 2 listed examples of designed molecules in these overlapping regions, showing their 2D UMAP coordinates, scaffold structures, representative compounds from PubChem, and FragLlama-designed molecules. This demonstrates that FragLlama’s ability to design drug-like molecules is enhanced when provided with comprehensive input data, including knowledge of the target protein, its known inhibitors, their bioassay results, and IC50 values, especially when more active compounds are incorporated into the fine-tuning process.

**Figure 13:**
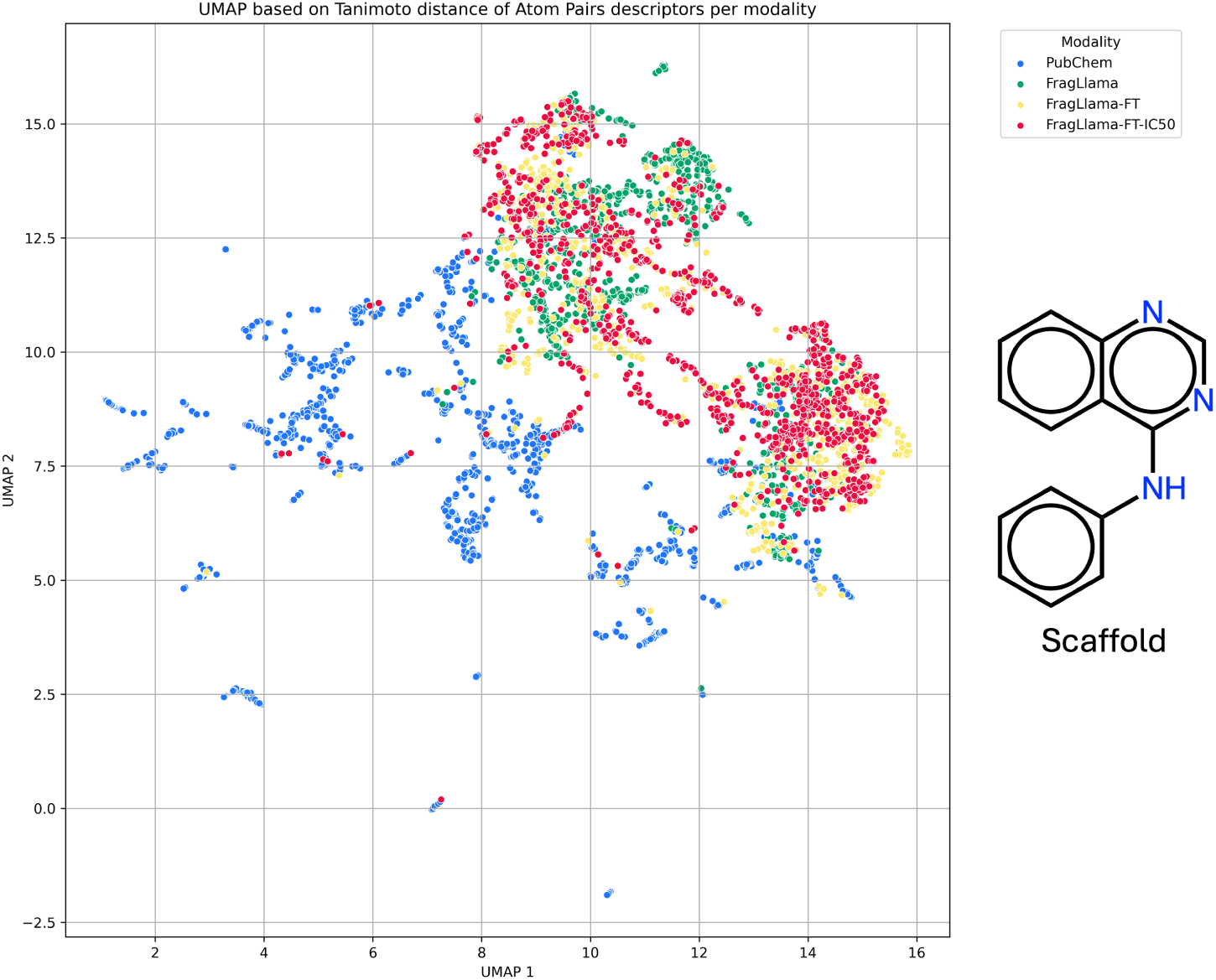
Chemical space distribution visualized using Uniform Manifold Approximation and Projection (UMAP), based on Tanimoto distances^53^ calculated from Atom-pair finger-prints.^52^ The plot compares three groups of molecules: 1) fine-tuned FragLlama designed molecules with both EGFR inhibitors and IC50 special tokens (red); 2) fine tuned FragLlama-designed molecules with only EGFR inhibitors (yellow); 3) non-fine-tuned FragLlama-designed molecules, alongside EGFR inhibitors from PubChem with active results in bioassays and valid IC50 values (blue).

**Table 2:** List of overlapping regions between fine-tuned FragLlama-designed molecules (trained with EGFR inhibitors and IC50 special tokens) and EGFR inhibitors from Pub-Chem. Each region includes 2D UMAP coordinates, scaffold structures, representative Pub-Chem compounds, and FragLlama-designed molecules.

| Region | Scaffold | Example compound in PubChem | FragLlama designed molecule |
| --- | --- | --- | --- |
| #1<br>(7.3, 0.2) |  |  |  |
| #2<br>(5.9, 11.0) |  |  |  |
| #3<br>(6.7, 7.8) |  |  |  |
| #4<br>(5.1, 7.6) |  |  |  |
| #5<br>(5.4, 8.2) |  |  |  |
| #6<br>(4.3, 7.7) |  |  |  |
| #7<br>(9.6, 8.4) |  |  |  |
| #8<br>(7.7, 12.2) |  |  |  |

## Conclusions

In conclusion, the core innovation of FragLlama lies in transforming traditional text token prediction into the prediction of molecular fragments, allowing the model to capture localized chemical features and common patterns in molecular structures. By progressively predicting and adding new fragments, FragLlama generates complete molecular structures while ensuring chemical reasonability and enhancing structural diversity. This fragment-level tokenization method bridges the gap between chemical structural representation and language model capabilities, improving the performance and applicability of language models in molecular design. By training tokens representing molecular fragments from scratch and leveraging a decoder-only architecture, FragLlama learns long-range dependencies that adhere to complex structural rules. The model demonstrates significant potential in drug discovery, by generating expert-level molecular designs and exploring new target-relevant chemical spaces.

## Acknowledgement

We thank Bryan Barker and Ruzhu Chen at Oracle Cloud for their expertise in high-performance computing infrastructure and technical assistance, and Prof. Jin Wang at Baylor College of Medicine for valuable discussions on molecular design case studies.

